# A computational approach for generating smooth estimates of motor unit discharge rates and visualizing population discharge characteristics

**DOI:** 10.1101/2021.08.09.455721

**Authors:** James. A. Beauchamp, Obaid U Khurram, Julius P.A. Dewald, Charles J. Heckman, Gregory E P Pearcey

## Abstract

**Objective:** Successive improvements in high density surface electromyography and decomposition techniques have facilitated an increasing yield in decomposed motor unit (MU) spike times. Though these advancements enhance the generalizability of findings and promote the application of MU discharge characteristics to inform the neural control of motor output, limitations remain. Specifically, 1) common approaches for generating smooth estimates of MU discharge rates introduce artifacts in quantification, which may bias findings, and 2) discharge characteristics of large MU populations are often difficult to visualize.

**Approach:** In the present study, we propose support vector regression (SVR) as an improved approach for generating continuous estimates of discharge rate and compare the fit characteristics of SVR to traditionally used methods, including Hanning window filtering and polynomial regression. Furthermore, we introduce ensembles as a method to visualize the discharge characteristics of large MU populations. We define ensembles as the average discharge profile of a subpopulation of MUs, composed of a time normalized ensemble average of all units within this subpopulation. Analysis was conducted with MUs decomposed from the tibialis anterior (N = 2128), medial gastrocnemius (N = 2673), and soleus (N = 1190) during isometric plantarflexion and dorsiflexion contractions.

**Main Result:** Compared to traditional approaches, we found SVR to alleviate commonly observed inaccuracies and produce significantly less absolute fit error in the initial phase of MU discharge and throughout the entire duration of discharge. Additionally, we found the visualization of MU populations as ensembles to intuitively represent population discharge characteristics with appropriate accuracy for visualization.

**Significance:** The results and methods outlined here provide an improved method for generating smooth estimates of MU discharge rate with SVR and present a unique approach to visualizing MU populations with ensembles. In combination, the use of SVR and generation of ensembles represent an efficient method for rendering population discharge characteristics.

## INTRODUCTION

Following its wide adoption into research, the electromyographic signal has been increasingly realized as a rich source of information regarding supraspinal and spinal mediated mechanisms for motor control. Specifically, given the tight coupling of action potentials (1:1 discharge) between spinal motor neurons and the muscle fibers that they innervate, collectively termed the motor unit, electric potentials recorded by both intramuscular and surface electromyography (EMG) function as a unique window into the central nervous system (Heckman & Enoka, 2012; Johnson, Thompson, Tysseling, Powers, & Heckman, 2017). Indeed, EMG signals are comprised of the superimposed action potentials of many motor units (MUs), which allows for individual MU discharge instances to be estimated with decomposition techniques. (De Luca, Adam, Wotiz, Gilmore, & Nawab, 2006; Farina, Holobar, Merletti, & Enoka, 2010; Holobar, Minetto, & Farina, 2014; Nawab, Chang, & De Luca, 2010; Rau & Disselhorst-Klug, 1997). These estimated MU discharge instances can then be used to characterize central nervous system function and garner insights into both healthy and pathological motor control, an approach bolstered through advancements in high density surface EMG (HD-sEMG) approaches (Kallenberg & Hermens, 2009; Li et al., 2015; Murphy et al., 2018).

Though the use of HD-sEMG and recent improvements in decomposition approaches have facilitated the application of estimated MU discharge characteristics to inform physiological understanding, its adoption into common practice and large scale application have introduced potential pit-falls in data estimation and visualization. Specifically, successive improvements in decomposition algorithms, HD-sEMG electrode arrays, and amplifier technology have continued to provide a greater yield in discriminated motor units. Though this increased MU yield has facilitated a greater confidence in the generalizability of findings and insight into population MU behavior, large increases in sample sizes often persuade researchers to reduce the dimensionality of their dataset through averaging, which fails to adequately account for variance, or be met with difficulties in accurately portraying the qualitative aspects of their data. This limitation will only magnify as an increasing amount of parameters are found to affect MU discharge patterns and is evident in recent papers from the field, where adequate qualitative representations of entire datasets are difficult to achieve and single choice trials are often displayed. A concise and informative methodology for portraying large sets of MU discharge profiles, or neuronal discharge in the general sense, would vastly improve the capability of researchers to relay their findings in a consistent and intuitive way.

More importantly, in addition to the difficulties of efficiently visualizing these increasingly large datasets, the process of extracting physiologically relevant metrics from decomposed MU discharge profiles is often variable amongst research groups and has considerable potential for biasing findings. Specifically, the process used to generate smooth estimates of MU discharge rates lacks an agreed upon computational method and has the potential to substantially influence frequently characterized outcome metrics. Commonly employed methods for obtaining smooth estimates of MU discharge rates include filtering of binary spike trains with a window function, such as the Hanning (Hann) window, or fitting instantaneous discharge rates with various degrees of lower order polynomial functions (Afsharipour et al., 2020; De Luca, LeFever, McCue, & Xenakis, 1982a; Gorassini, Yang, Siu, & Bennett, 2002; F. Negro & Farina, 2012). Though commonly used, filtering with the Hanning window introduces undesirable characteristics at the onset and offset of discharge, which biases the estimated recruitment and derecruitment discharge rates, and lower order polynomial functions can potentially remove relevant characteristics of MU discharge (i.e. over-smoothing). Indeed, a recent study demonstrated how fit method (Hanning, Gaussian, 5^th^-order polynomial) can affect estimates of persistent inward currents (PICs) generated by the paired MU analysis technique (i.e. ΔF), showing the edge effects of the Hanning window to bias motor unit recruitment and derecruitment estimates (Hassan et al., 2020). To minimize the introduction of biases in the data analysis pipeline, a method of generating smooth estimates of MU discharge rates that accurately represent the end conditions and more effectively balances the tradeoff between noise mitigation and retaining relevant discharge characteristics is necessary.

To address these problems, we 1) investigated support vector regression (SVR) as a more effective means of producing smooth continuous estimates of MU discharge rates and 2) propose that large populations of MUs be quantified and visualized in ensembles, or average traces of MU discharge rates for subpopulations of MUs separated by a metric of interest (e.g. torque at MU recruitment). Each ensemble represents the average behavior of motor units within a subpopulation and is composed of a time normalized estimate of discharge rate for each individual MU, generated through SVR. Support vector regression, with its ability to independently weight observations (e.g. MU recruitment and derecruitment) and tune hyperparameters to optimize fit, offers a level of control far superior to traditional fitting schemes (Alex J. Smola & Schölkopf, 2004; Vapnik, 1995). In specific, weighting the end conditions alleviates the biasing effects at recruitment and derecruitment introduced by the edge effects of the Hanning window while hyperparameter tuning tempers the unnecessary smoothing introduced by fitting with lower-order polynomial functions. We hypothesized that: 1) compared to the Hanning window and a 5^th^ and 6^th^ order polynomial, the capabilities inherent to SVR would facilitate a more accurate representation of estimated MU discharge rates, and 2) visualizing groups of MU discharge rates as ensembles would provide an intuitive method to convey findings in a compelling manner, where one figure can visually display the potential findings of an entire dataset.

## METHODS

### Dataset

#### Participants

Motor unit spike trains were obtained from multiple ongoing human subject studies. This included twenty-one young participants (F: 5, M: 16; Age: 26.4 ± 1.7) with no known neuromuscular, musculoskeletal, or cardiovascular impairments. All participants provided written and informed consent (Northwestern University Institutional Review Board STU00202964) in accordance with the Declaration of Helsinki.

#### Overview

Given that a primary goal of this effort was to provide data quantification and visualization methodologies for studies that employ estimates of smooth MU discharge rates, the contraction profile and muscles were chosen accordingly. Specifically, to assist in generalizability to future studies, a ramp contraction, consisting of a linear increase and subsequent decrease in effort, was chosen because it provides desirable MU recruitment spacing and is commonly used in the field (De Luca, LeFever, McCue, & Xenakis, 1982b; Farina et al., 2009; Kim, Wilson, Thompson, & Heckman, 2020; Orssatto et al., 2021; Oya, Riek, & Cresswell, 2009). Similarly, given frequent use of the lower limb in HD-sEMG studies, ramp contractions were generated through either ankle dorsiflexion or plantarflexion with grid electrodes fixed atop the skin overlying the tibialis anterior (TA), medial gastrocnemius (MG), and soleus (SOL) muscle bellies.

#### Experimental Setup

For each experimental session, participants were seated in a Biodex chair, with their left foot securely attached to a footplate fixed onto a Systems 4 Dynamometer (Biodex Medical Systems, Shirley, NY) such that the axis of rotation aligned with the center of rotation of the ankle joint. Throughout the session a participants’ hips were maintained at approximately 80 degrees of flexion, left knee at 20 degrees flexion, and left ankle at 10 degrees of plantarflexion with thigh and shoulder straps used to minimize movement. Target torque ramps and visual feedback (i.e. dorsiflexion or plantarflexion torque) were provided on a television screen via a custom Matlab interface (MATLAB (R2020b), The Mathworks Inc., Natick, MA). Torque about the ankle was filtered with a 125 ms moving average window before being provided as visual feedback to the participant. For subsequent analysis, raw torque signals were amplified (150 ×) and digitized (2048 Hz) using a 16-bit analog-to-digital converter (Quattrocento, OT Bioelettronica, Turin, IT) and lowpass filtered (50 Hz) with a fifth order Butterworth filter.

#### Experimental Protocol

Prior to commencement of ramp contractions, participants were asked to generate maximal voluntary isometric contractions of the plantarflexors and dorsiflexors, with 2 minutes of rest separating contractions. At least two contractions were performed, and repeated until the peak torque within the last contraction was no larger than 5% of the previous contraction. We then used the maximum voluntary torque (MVT) achieved during these contractions to normalize all subsequent ramp contractions. Ramp contractions started from rest and consisted of a 10 second linear increase to 30% MVT and a 10 second decrease back to rest (i.e. 3% MVT/s rise and decay speeds). To mitigate learning effects and ensure smooth contractions, participants completed a minimum of 6 dorsiflexion and plantarflexion practice ramps. Following practice trials, each experimental session consisted of 4-12 ramp contractions for each dorsiflexion and plantarflexion, conducted in random order.

#### High Density Surface EMG (HD-sEMG)

HD-sEMG was collected via 64 channel electrode grids (GR08MM1305, OT Bioelettronica, Turin, IT) placed atop the skin with adhesive foam (KITAD064, OT Bioelettronica, Turin, IT) overlying the TA, MG, and SOL muscle bellies. The location of the muscles were identified via palpation by a clinical exercise physiologist. Prior to electrode placement, the left leg was shaved and the skin overlying the muscles was abraded with abrasive paste and cleaned with isopropyl alcohol. Two Ag/AgCl ground electrodes were placed bilaterally on the right and left patella and a moist band electrode was placed around the right ankle. HD-sEMG signals were acquired with differential amplification (150 x), digitized (2048 Hz), and bandpass filtered (10-900 Hz) using a 16-bit analog-to-digital converter (Quattrocento, OT Bioelettronica, Turin, IT).

#### Motor Unit Decomposition

In preparation for decomposition, all surface EMG channels were bandpass filtered at 20–500 Hz (second-order, Butterworth) and visually inspected to remove channels with substantial artifacts, noise, or saturation of the A/D board (typically 2-3 channels). The remaining EMG channels were decomposed into individual MU spike trains using convolutive blind source separation and successive sparse deflation improvements (Martinez-Valdes et al., 2017; Francesco Negro, Muceli, Castronovo, Holobar, & Farina, 2016). The silhouette threshold for decomposition was set to 0.87. To improve decomposition accuracy and correct spikes that indicated non-physiological MU discharge, experienced investigators conducted manual editing of the spike trains. Specifically, automatic decomposition results were improved through iteratively re-estimating the spike train and correcting for missed spikes or substantial deviations in the discharge profile (Boccia, Martinez-Valdes, Negro, Rainoldi, & Falla, 2019; Del Vecchio et al., 2020; Hug et al., 2021).

### Computational Fitting Methods

To compare support vector regression (SVR) with commonly employed methods, smooth MU discharge rates were generated with the following computational approaches. For Hanning (Hann) window filtering, analysis began with decomposed binary spike trains, whereas SVR and polynomial regression was initiated with discrete estimates of instantaneous discharge rate. To obtain these discrete values, estimated MU discharge times were obtained from decomposed MU spike trains and used to quantify the inter-spike interval (ISI), or the time between each consecutive spike. A discrete estimate of instantaneous discharge rate was then calculated as the reciprocal of the time series ISI for each MU. For all trials, any MU which failed to sustain a minimum of 10 consecutive discharges was removed from analysis. This resulted in a total MU Yield of 2128 for TA, 2673 for MG, and 1190 for SOL.

#### Hanning (Hann) Window Filtering

The use of the Hanning window to generate smooth discharge rate estimates from decomposed motor unit spike trains has been a popular computational approach within the field (De Luca et al., 1982a). To obtain these estimates, a binarized motor unit spike train is filtered with a Hanning window of pre-specified length. We have chosen to employ a Hanning window with length equivalent to 1 s in duration, though windows of various durations have previously been employed (De Luca et al., 1982a; Hassan et al., 2020; F. Negro & Farina, 2012).

#### Polynomial Regression

Fitting instantaneous discharge rates with a polynomial function is becoming a common practice amongst the field to provide smooth discharge rate estimates, with a 5^th^ order polynomial commonly used (Afsharipour et al., 2020; Gorassini et al., 2002). To compare SVR and polynomial regression, we have chosen to use a 5^th^ and 6^th^ order polynomial function to fit instantaneous discharge rates. Given that a 5^th^ order polynomial necessitates opposite end conditions, an even degree polynomial should theoretically better represent the discharge profiles observed during isometric ramp contractions. Furthermore, an even degree polynomial greater than a 5^th^ order function is likely more desirable (i.e. 6^th^ not 4^th^ order), given the known smoothing properties. Polynomial regression of the instantaneous discharge rates was accomplished through employing least squares with the time vector centered at zero and scaled to one standard deviation. The polynomial coefficients produced by this operation were then used to generate smooth estimates of discharge rate along a prediction time vector from MU recruitment to derecruitment sampled at 2048 Hz.

#### Support Vector Regression

Support vector regression was first introduced by Vapnik and colleagues, who outlined the application of traditional support vector machine classification to a regression problem (Drucker, Burges, Kaufman, Smola, & Vapnik, 1996; Vapnik, 1995). Like classification with support vector machines, support vector regression (SVR) employs much of the same principles, including use of kernels to represent data in a higher dimensional space, a hyperplane separating data points in this higher dimensional space, and a margin about this hyperplane. In depth discussions on SVR and its algorithmic implementation can be found elsewhere and will not be included here (Cristianini & Shawe-Taylor, 2000; Alex J Smola & Schölkopf, 1998; Alex J. Smola & Schölkopf, 2004). For ease of translation across research groups, we implemented SVR with Matlab’s inbuilt function *fitrsvm* to train an SVR model with L1 soft-margin minimization (MATLAB (R2020b), The Mathworks Inc., Natick, MA). For each MU, training data included the instantaneous discharge rate estimates and corresponding time instances. Smooth estimates of discharge rate were then generated using Matlab’s inbuilt *predict* function to generate an estimated discharge rate along a prediction time vector from MU recruitment to derecruitment sampled at 2048 Hz (MATLAB (R2020b), The Mathworks Inc., Natick, MA).

Support vector regression contains various parameters that can be tuned to optimize fitting characteristics. For our purposes, this included the kernel that is employed, the kernel scale factor, epsilon, and the regularization parameter (Alex J. Smola & Schölkopf, 2004). The kernel and kernel scale factor define the function employed in expanding dimensionality and was chosen as a radial basis function. Epsilon defines one-half of the margin, or width about the hyperplane in which no penalty is assigned to the cost function. The regularization parameter indicates the penalty that is assigned to points outside this margin.

To account for the inherent differences in variability of discharge between muscles, we chose an epsilon value that was MU specific and scaled based on the discharge variability for that unit. Specifically, we chose an epsilon value equal to one-eleventh of the interquartile range of the discharge rate, which generates an approximate epsilon insensitive region (margin) of one quarter of a standard deviation. To ensure desirable end characteristics, the initial and final five discharge instances were weighted five times greater than the remainder of discharge instances for a given unit. To optimize the kernel scale and regularization parameters, we performed a grid search across a range of 0-1000 for both terms and used the values that most closely replicated the average sum of squared error achieved with the 1 s Hanning window throughout the middle 80% of discharge. That is, the error seen at the first and last 10% of discharge was not considered given the known edge effects introduced by filtering with the Hanning window. The 1s Hanning window was chosen as a comparator, given that this fit is generally believed to retain and accurately portray the relevant characteristics of MU discharge throughout the *middle* portion of discharge. This produced a regularization parameter of 370 and a kernel scale factor of 1.6.

### Ensembles

An overview of the construction of ensembles can be seen in Figure 1. For each muscle, all discriminated MUs across participants (TA: 2128, MG: 2673, SOL: 1190) were separated into ten equally spaced bins based upon the percent of Maximum Voluntary Torque (MVT) that a MU was recruited at (i.e. 3% MVT increments for 30% MVT ramps). The MUs within each of these 3% MVT bins were then fit with the three computational fitting methods. For each fit method, we utilized a time normalization procedure to generate MUs of a pre-specified length within each ensemble. For SVR and polynomial regression, we adjusted the sampling rate of the prediction vector such that smooth discharge rate estimates within an ensemble were vectors of equal length, equivalent to the length of the time vector from average MU recruitment to derecruitment sampled at 2048 Hz. Similarly, for Hanning window filtering, the smooth discharge rate estimates generated by the Hanning window were resampled using linear interpolation to generate vectors of identical length within a given ensemble. Following this normalization, we then quantified the ensemble average of all MU fits within each 3% MVT cohort and mapped these traces from the average recruitment to derecruitment instance of each group to generate the quantized ensemble traces. This was done such that each ensemble trace represented a true “average”, with the ensemble discharge rate traces representing the average discharge profile from MU recruitment to MU derecruitment for all units within that cohort.

**Figure 1:**
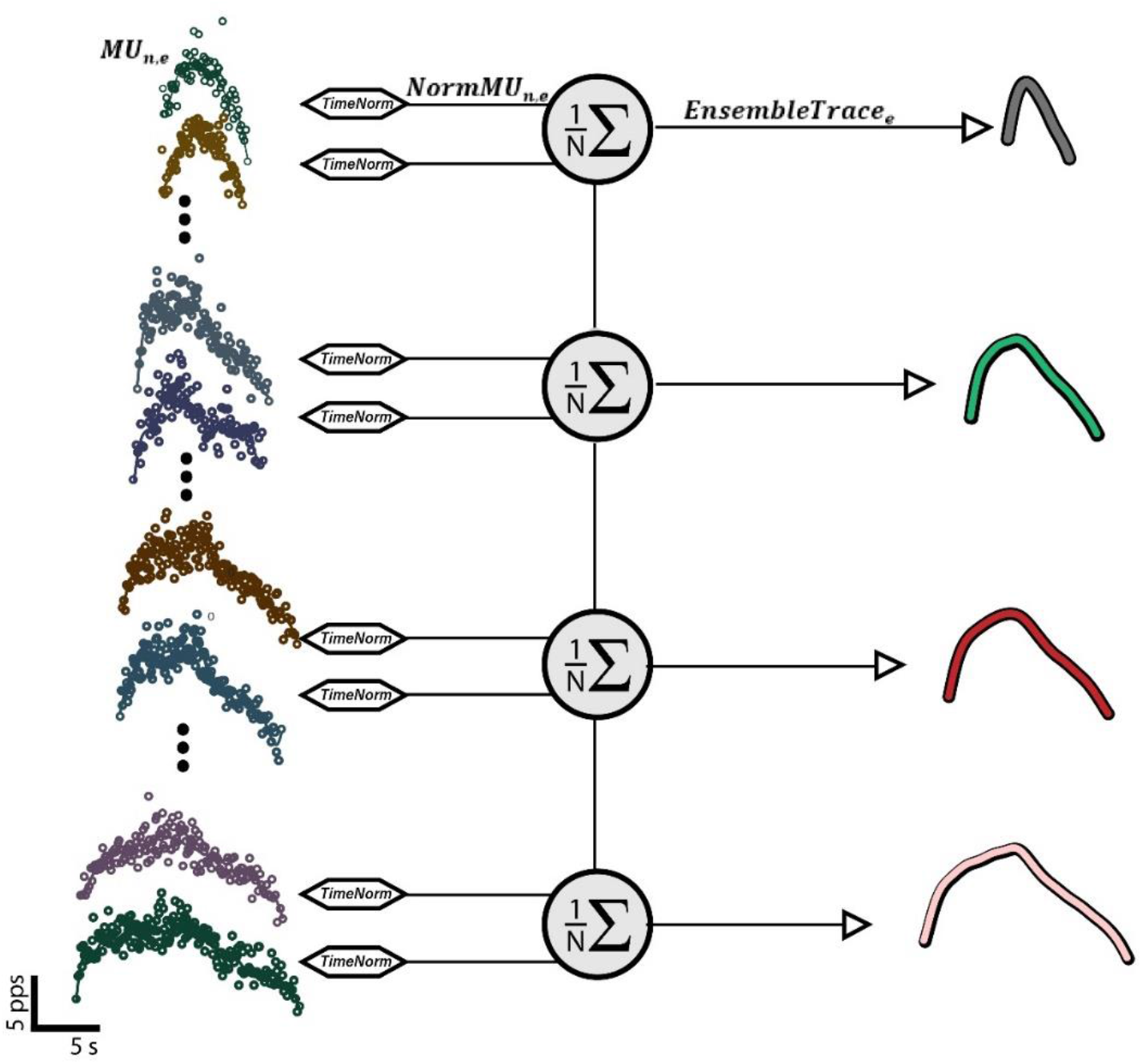
Overview of ensemble construction. Populations of estimated motor unit discharge profiles were subdivided into cohorts based upon a metric of interest (four groups on left), filtered or fit with an estimating function with the time axis (x-axis) normalized such that motor units within a subdivision align from onset to offset, and ensemble averaged (four traces on right). (pps: pulses per second; s: second)

To further illuminate the time normalization process, Figure 2 shows the SVR estimates for all TA MUs (N = 2128) separated into ten ensemble cohorts both before (Figure 2A; non-normalized) and after (Figure 2B; normalized) normalization. In this figure, the quantized SVR ensemble traces (Figure 2A: black traces) can be seen to unsurprisingly represent the average onset and offset of discharge across MUs, given that this is how they were defined. Furthermore, in the time normalized traces, representation of MUs on an identical timescale from recruitment to derecruitment can be observed. This allows for the ensemble trace to represent the true average shape of MUs within each ensemble, with distinct modes of discharge mapped from onset to offset. The ability of these ensemble traces to represent the average discharge profile of each MU cohort can be seen with the black traces overlying the non-normalized fits.

**Figure 2:**
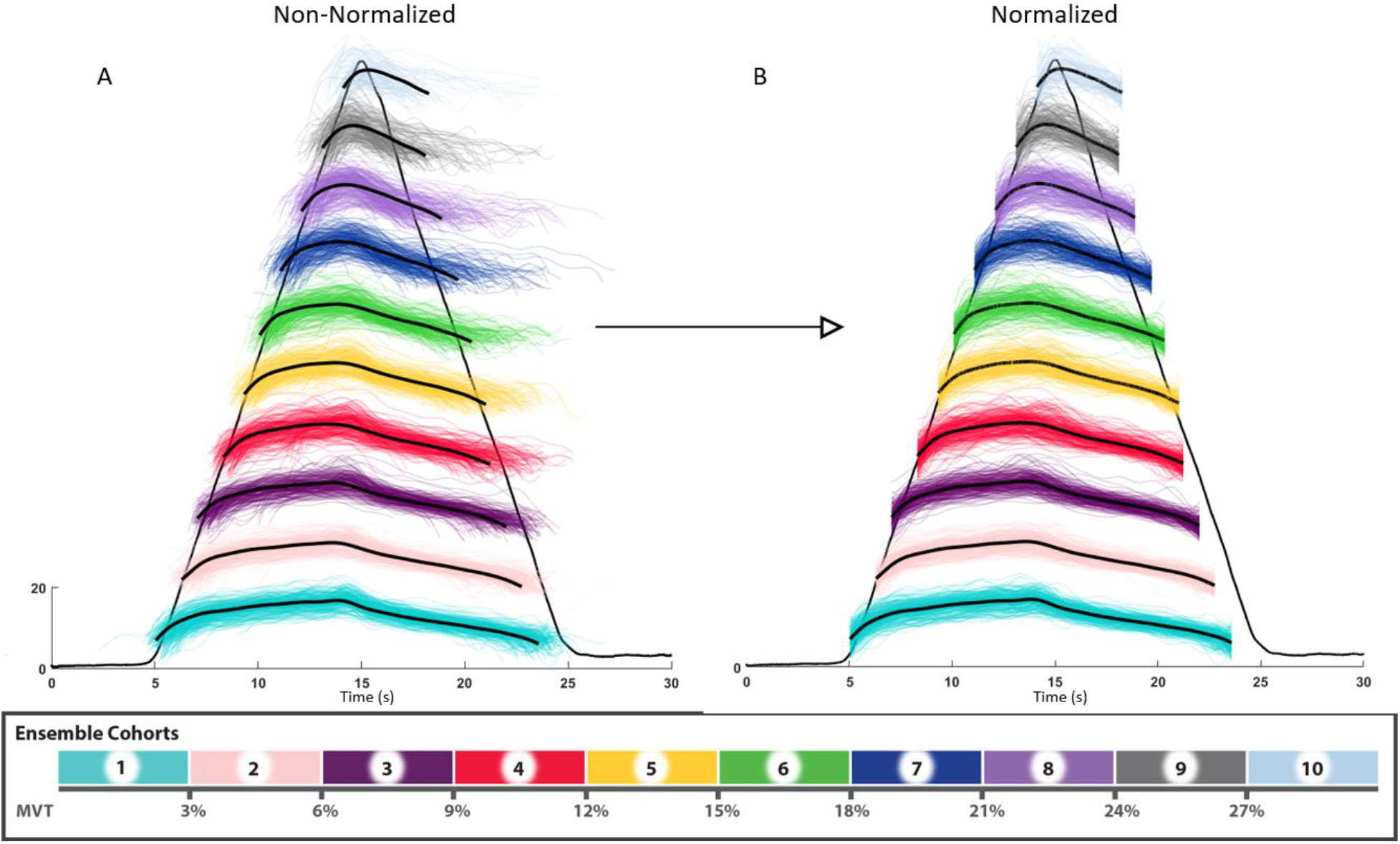
Normalization in ensemble construction. Shown above is the ensemble construction process for all decomposed TA motor units (N = 2128). Units are first separated into ten cohorts based upon their recruitment threshold, with the color bar indicating the recruitment range of this group normalized to individual participants’ MVT and the black triangular trace representing the average torque across all trials and participants. Smoothed estimates of MU discharge rate are then created with support vector regression (A), and projected onto time vectors of identical length (B). These normalized estimates are then ensemble averaged to generate the overlying black traces. These ensemble traces in black are shown overlying the normalized and non-normalized estimates for comparison. (pps: pulses per second; s: second)

### Fit Comparison & Ensemble Accuracy

To compare the accuracy of the three computational fitting methods we quantified the residual error, or absolute deviation between the continuous and discrete discharge rate estimates, for each individual MU. This was conducted to provide insight into the type of bias introduced by each fit, spanning from MU recruitment to derecruitment. To observe the qualitative effects of the various fitting schemes, ensemble figures were constructed for each fit method.

#### Simulations

To characterize the accuracy of representing populations of MUs as ensembles, we utilized a Monte Carlo type simulation. Specifically, we treated the process of creating ensemble traces as a transfer function, decomposed MU spike trains as inputs, and used outcome metrics for each individual MU and ensemble trace of discharge rate at recruitment, discharge rate at derecruitment, peak discharge rate, time to peak discharge rate, and ΔF.

For each iteration, we separated the total MU dataset for the TA into ensemble groups based upon their torque at recruitment, as before, and iteratively resampled two-thirds of this population. This was conducted for 100 iterations, with the average population outcome metrics for all units within each ensemble and the outcome metrics for the ensemble trace used to generate estimated distributions for each metric of interest. The difference between the estimated distribution of the sampled MU populations and ensemble traces was then used to garner insight into the ability of the ensemble traces to capture the characteristics of the populations of MUs within them.

Discharge rate at recruitment and derecruitment were calculated as the discharge rate of the first and last instance of the smooth SVR fits, respectively. Peak discharge was calculated as the maximum discharge rate of each MU SVR fit or ensemble trace, with time to peak calculated as the time between MU recruitment and this peak value.

ΔF is a commonly employed metric used estimate the magnitude of PICs and represents the discharge hysteresis of a higher threshold MU with respect to a lower threshold unit. To quantify ΔF, we employed a paired MU analysis technique such that ΔF for a given MU (test unit) represented the change in discharge rate of a lower threshold unit (reporter unit) between the recruitment and derecruitment of this test unit. This was conducted for every possible combination of MU pairs within a trial where the reporter unit exhibited sustained discharge throughout the test units recruitment and derecruitment. To account for the pairing of a test unit with multiple reporter units, ΔF for a test unit was calculated as the average change in discharge rate across all possible reporter unit pairs. To allow for full activation of the PIC in the reporter unit, we excluded any pairs with recruitment time differences <1 s (Bennett, Li, Harvey, & Gorassini, 2001; Hassan et al., 2020; Powers, Nardelli, & Cope, 2008). Additionally, to avoid saturated reporter units, we excluded test unit-reporter unit pairs in which the reporter unit discharge range was < 0.5 pps while the test unit was active (Stephenson & Maluf, 2011). Furthermore, we only included test unit-reporter unit pairs with rate-rate correlations of r^2^ > 0.7 to ensure that MU pairs likely received common synaptic drive (Gorassini et al., 2002; Udina, D’Amico, Bergquist, & Gorassini, 2010; Wilson, Thompson, Miller, & Heckman, 2015).

To quantify ΔF for a given ensemble, we conducted a similar process treating each ensemble as either a test or reporter ensemble. Specifically, all ensemble traces of a lower recruitment torque than a given ensemble were used as reporter ensembles. ΔF for a test ensemble was then calculated as the average change in discharge rate across all possible reporter ensembles from test ensemble recruitment to derecruitment.

### Statistical Approach

To determine significant differences in fit error between the fitting methods, we employed linear mixed effects models with either average absolute fit error across the first five discharges of a MU or the entire duration of MU discharge as a dependent variable, fixed effects of torque recruitment cohort (ensemble), muscle, and fit method, and random effects of participant and trial nested within participant. P-values were obtained by likelihood ratio tests of the full model with the effect in question against the model without the effect in question. For main effects, this included their subsequent interaction terms.

To test for statistically significant differences in discharge characteristics of interest between SVR ensembles and the MU population characteristics, we used data generated with the Monte Carlo simulation. Specifically, we employed linear models with dependent variables of the ensemble and sample estimates for each outcome metrics of interest. For simplicity we only analyzed the TA data set and included a total of 1000 observations for each method (ensemble or sample estimate) and outcome metric, or 100 iterations for each of the ten ensemble cohorts. As fixed effects we included torque recruitment cohort (ensemble), whether the estimate was from the ensemble trace or sample population average, and the interaction between these factors.

All statistical analysis was performed with R (R Core Team, 2021). Mixed model analysis was achieved via the *lme4* (Bats, Maechler, Bolker, & Walker, 2015) package. To ensure the validity of model fitting, the assumptions of linearity and normal, homoscedastic residual distributions were confirmed. Estimated marginal means were employed in pairwise post-hoc testing and achieved with the *emmeans* package (Lenth, 2021). Significance was set at α = 0.05 and pairwise and multiple comparisons were corrected using Tukey’s corrections for multiple comparison.

## RESULTS

### Fit Method Comparison

A randomly chosen MU from the MG muscle during a single plantarflexion trial can be seen in Figure 3 with each fit method applied. Qualitatively, polynomial regression has produced estimates that follow the general trend of the MU discharge profile but fail to portray key attributes of discharge associated with the activation PICs. This includes an initial high gain phase (PIC activation or acceleration of discharge) followed by a gain attenuated phase (PIC saturation or post-acceleration discharge rate saturation) (Heckman & Enoka, 2012). Additionally, on the descending portion of the ramp (~16-18 s) an abrupt decrease in torque occurs with a corresponding decrease in discharge rate that is not captured by either polynomial fit. Support vector regression and Hanning window filtering, with the employed parameters, do appear to follow the discharge profile during this decrease in torque as well as highlight the expected features introduced by PICs. That said, while these attributes are captured by the Hanning window filtering approach, this fit appears to underestimate the initial discharge rate of 5.77 pps by a sizable margin (Hann: 3.94 pps, Poly-5: 7.41 pps, Poly-6: 6.72 pps, SVR: 5.73 pps).

**Figure 3:**
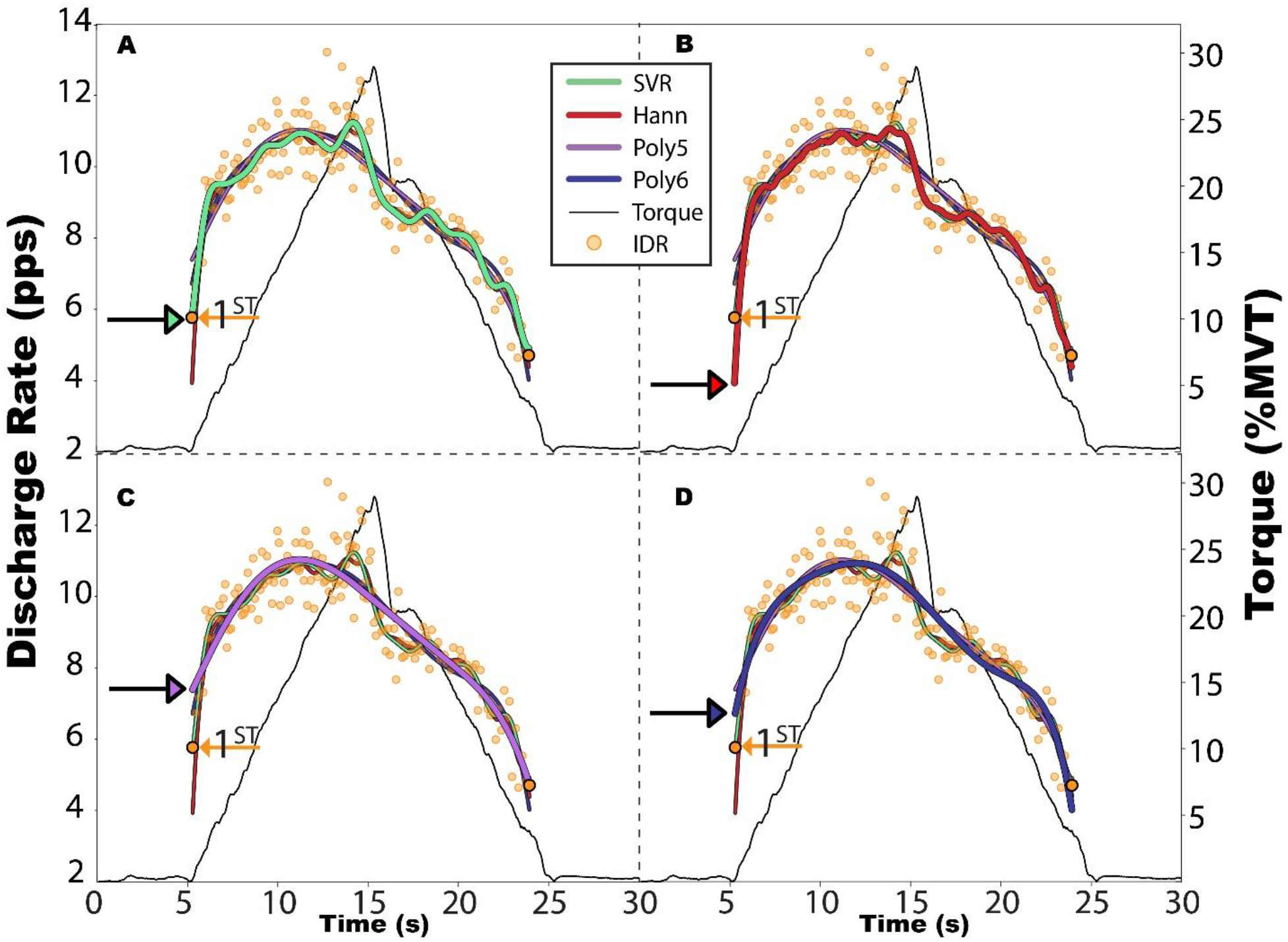
Effects of fit method on smoothed estimates of discharge for a single motor unit. A single decomposed medial gastrocnemius (MG) motor unit from a plantarflexion trial is shown with its estimated instantaneous discharge rate (IDR) in pulses per second (pps) and orange. The discharge rate at recruitment and derecruitment are shown outlined in black. The smooth estimates of discharge rate generated through each investigated approach are shown in solid lines. This includes A) support vector regression (SVR), B) Hanning (Hann) window filtering, and C-D) polynomial regression with either a 5^th^ or 6^th^ degree polynomial. Plantarflexion torque about the ankle is shown in black, normalized to maximum voluntary torque (MVT).

To highlight the fitting characteristic of each computational approach, Figure 4 shows the absolute fit error for each method as a function of unit duration. As is observed, the fit error is unequally distributed across the discharge profile for each fit method, with the various fitting schemes diverging at the onset and offset of discharge and the polynomial regression fits systemically higher across the unit duration. The inset within Figure 4 accentuates this divergence at onset, showing the absolute fit error averaged across all units for the first five instances of all MUs discharge.

**Figure 4:**
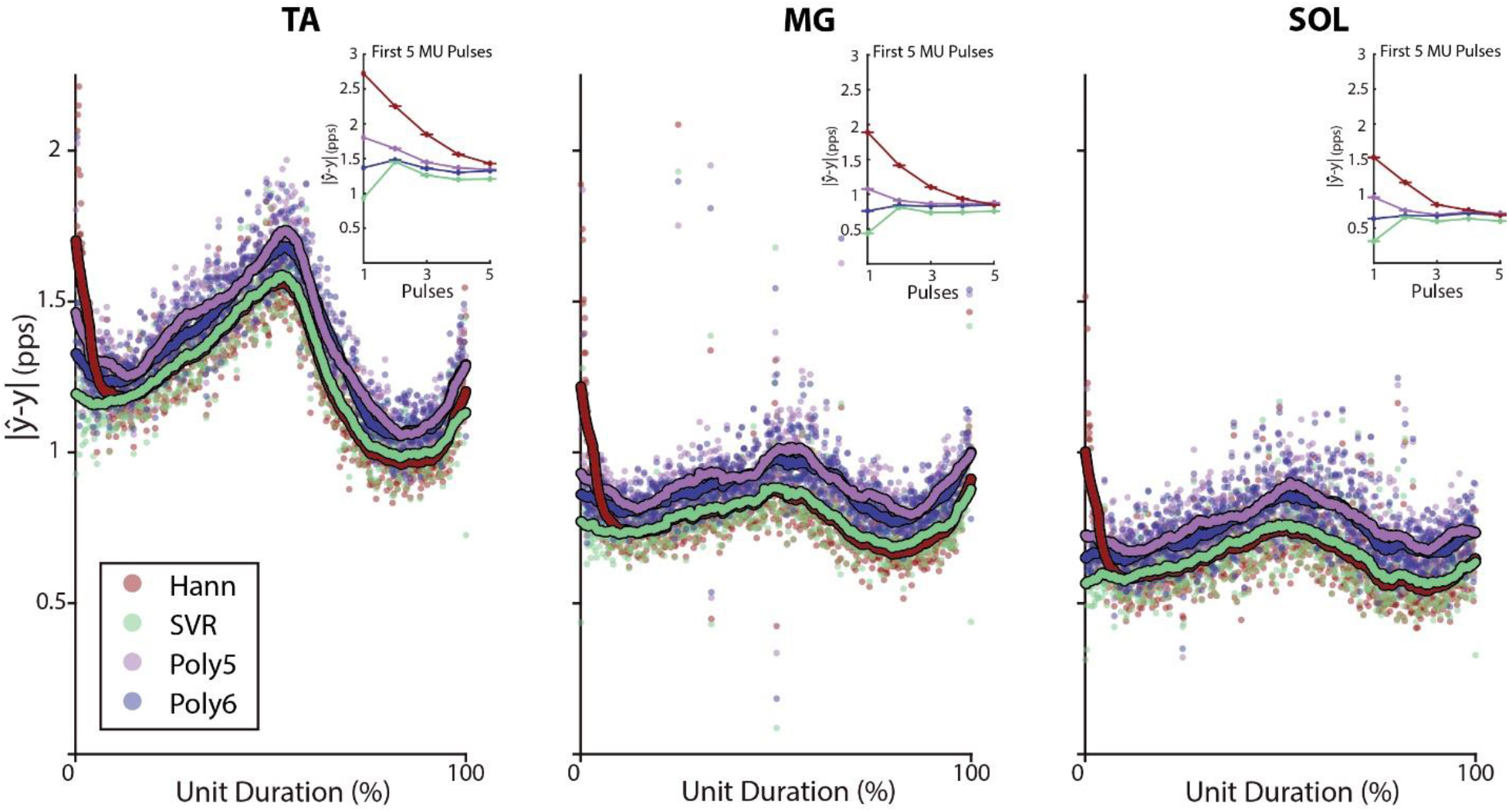
Absolute fit error for each fit method across motor unit duration. The absolute difference between the estimated discharge rate (ŷ) and instantaneous discharge rate (y) is shown as a function of motor unit duration from recruitment (0%) to derecruitment (100%) for each fit method in pulses per second (pps). Fit methods included support vector regression (SVR), Hanning (Hann) window filtering, and polynomial regression with either a 5^th^ or 6^th^ degree polynomial. Colored dots indicate individual discharge instances and the solid lines represent a moving average with width corresponding to a 6% unit duration. The inset displays the absolute error plot as a function of the number of discharges from recruitment, for the first five discharges for all motor units (avg. ± SE). This is shown for the tibialis anterior (TA, N = 2128), medial gastrocnemius (MG, N = 2673), and soleus (SOL, N = 1190).

To further investigate this divergence of fitting schemes, the average fit error for the first five discharges for each MU can be seen in the top row of Figure 5A with a corresponding probability density function. We used a linear mixed model and maximum likelihood estimation to predict this fit error within the first five discharges of a MU (Absolute Error ~ FitMethod*Muscle*Ensemble + (1|PID:Trial)), showing fit method (χ^2^(90) = 4082.4, p < 0.001), ensemble cohort (χ^2^(108) = 1959.5, p < 0.001), and muscle (χ^2^(80) = 1140.7, p < 0.001) to be significant predictors of this fit error. Additionally, interactions between fit method and ensemble (χ^2^(81) = 394.2, p < 0.001), fit method and muscle (χ^2^(60) = 194.6, p < 0.001), and muscle and ensemble were observed (χ^2^(72) = 548.3, p < 0.001). Across all muscles and ensemble cohorts, a main effect comparison of marginal means yields an estimated decrease in absolute error with SVR when compared to Hanning (0.584 pps; 95%CI: [0.554, 0.614]), 5^th^ degree polynomial (0.225 pps; 95%CI: [0.195, 0.256]), and 6^th^ degree polynomial (0.114 pps; 95%CI: [0.083, 0.144]). Significant decreases in absolute error with SVR were also observed when separated by muscle for the difference between SVR and Hanning (TA: 0.758 pps [0.714, 0.802]; MG: 0.564 pps [0.521, 0.608]; SOL: 0.430 pps [0.363, 0.496]), SVR and 5^th^ degree polynomial (TA: 0.297 pps [0.252, 0.341]; MG: 0.197 pps [0.154, 0.241]; SOL: 0.182 pps [0.116, 0.249]), and SVR and 6^th^ degree polynomial (TA: 0.142 pps [0.098, 0.187]; MG: 0.098 pps [0.055, 0.142]; SOL: 0.100 pps [0.034, 0.167]).

**Figure 5:**
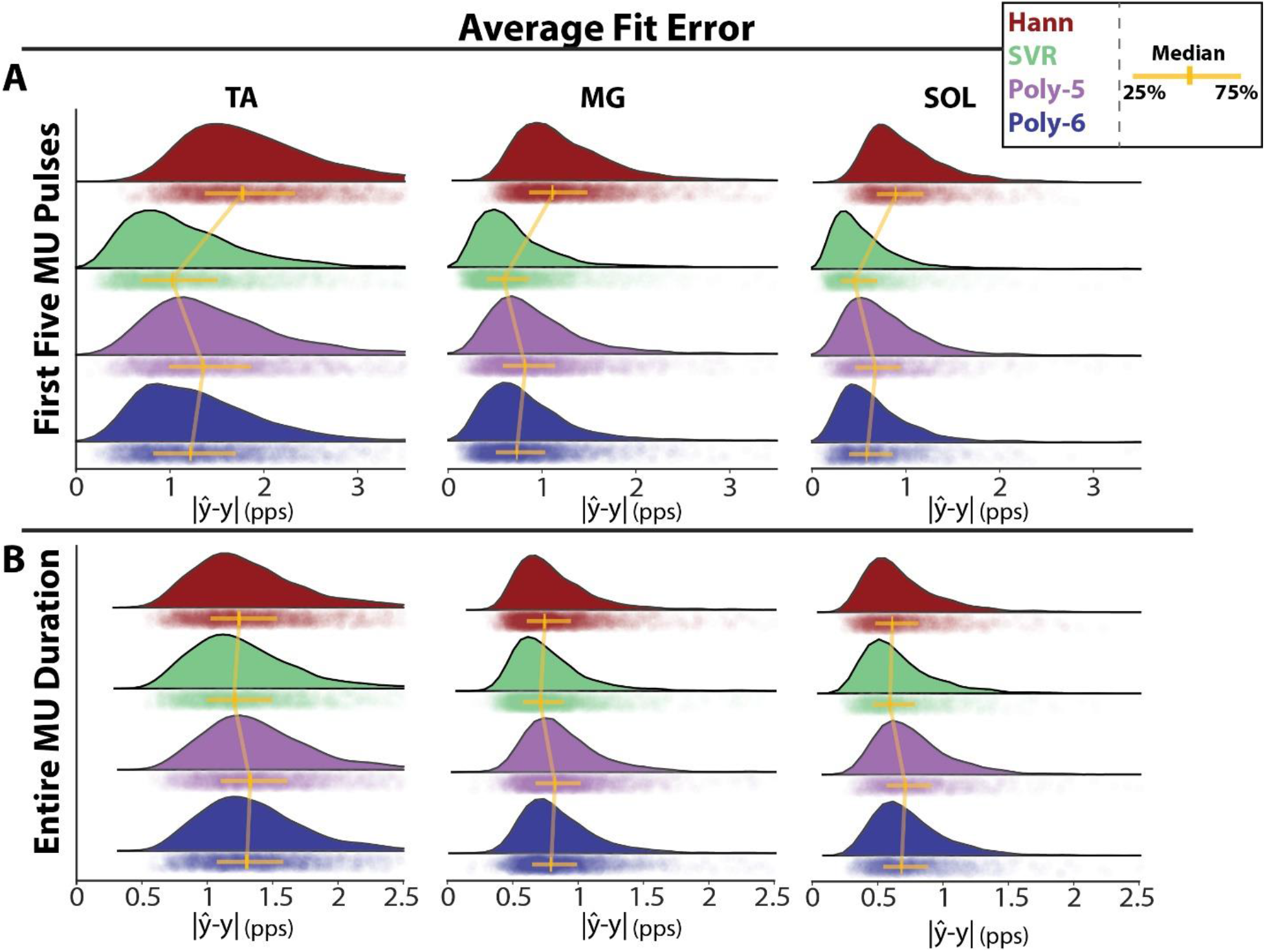
Average absolute error across the fit methods. The average absolute difference between the estimated discharge rate (ŷ) and instantaneous discharge rate (y) is shown for each fit method in pulses per second (pps) for the first five motor unit discharges (A) and entire duration of discharge (B). Each data point represents an individual motor unit with the corresponding probability density generated by a gaussian kernel. Individual colors represent the various fit methods and include support vector regression (SVR), Hanning (Hann) window filtering, and polynomial regression with either a 5th or 6th degree polynomial. This is shown for the tibialis anterior (TA, N = 2128, left), medial gastrocnemius (MG, N = 2673, middle), and soleus (SOL, N = 1190, right).

Using average fit error across the entire MU discharge duration (Figure 5B) as the dependent variable in a linear mixed effects model (Absolute Error ~ FitMethod*Muscle*Ensemble + (1|PID:Trial)), we found fit method (χ^2^(90) = 637.1, p < 0.001), muscle (χ^2^(80) = 929.9, p < 0.001), and ensemble (χ^2^(108) = 2605.7, p < 0.001) to be significant predictors of fit error. Furthermore, we observed an interaction between fit method and ensemble (χ^2^(81) = 205.0, p < 0.001) as well as muscle and ensemble (χ^2^(72) = 397.0, p < 0.001) with a non-significant fit method and muscle interaction (χ^2^(60) = 15.0, p ~=1). Across all muscles and ensemble cohorts, marginal means for the absolute error are estimated as 0.949 pps (95%CI: [0.915, 0.982]) for SVR, 1.052 pps (95%CI: [1.019, 1.086]) for the 5^th^ degree polynomials, 1.021 pps (95%CI: [0.987, 1.054]) for the 6^th^ degree polynomials, and 0.990 pps (95%CI: [0.957, 1.024]) for the Hanning window. When separated by muscle, we observed significant decreases in marginal means for absolute error with SVR when comparing the difference between SVR and 5^th^ degree polynomial (TA: 0.106 pps [0.080, 0.132]; MG: 0.103 pps [0.077, 0.128]; SOL: 0.102 pps [0.063, 0.141]), SVR and 6^th^ degree polynomial (TA: 0.074 pps [0.048, 0.101]; MG: 0.068 pps [0.042, 0.094]; SOL: 0.075 pps [0.035, 0.114]), and SVR and Hanning (TA: 0.049 pps [0.023, 0.075]; MG: 0.056 pps [0.030, 0.081]; SOL: 0.021 pps [0.019, 0.060]).

To illustrate the impact of each fit method on ensemble construction, we display the ensemble traces for each of the four fit methods applied to all of the TA MUs in Figure 6. The over-smoothing created through polynomial regression and the biased end conditions of Hanning window filtering are highly evident and correspond to the fit errors that were observed for each method (Figures 4 & 5).

**Figure 6:**
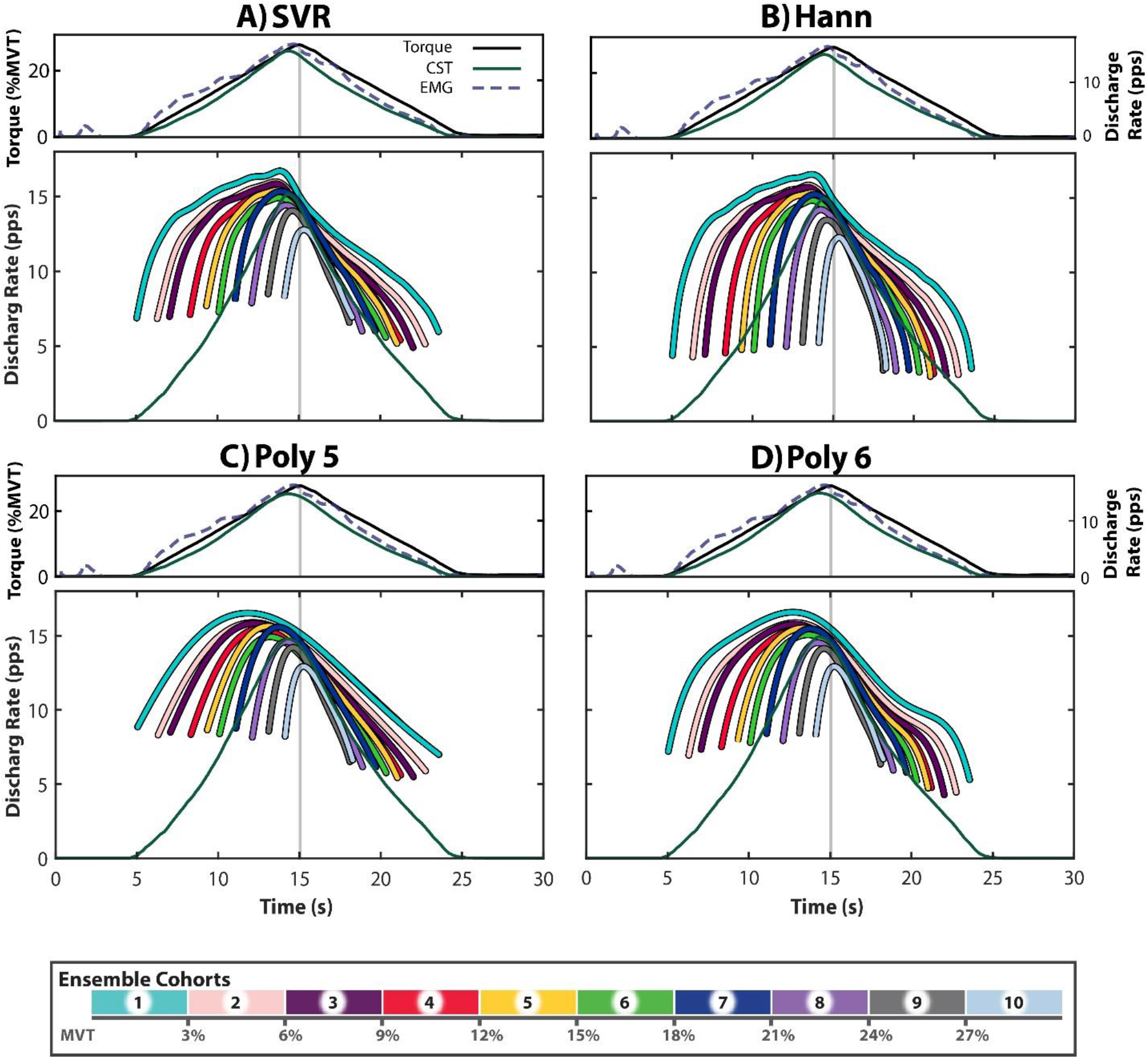
Ensembles for each fit method. Each quadrant represents one of the investigated fitting schemes and corresponds to A) support vector regression (SVR), B) Hanning (Hann) window filtering, and C-D) polynomial regression with either a 5^th^ or 6^th^ degree polynomial. Within each quadrant, the top plot depicts the average torque in black, the cumulative spike train (CST) across all units in green, and the average moving root mean squared EMG for all trials in dashed purple. Torque is shown as percent of maximum voluntary torque (MVT), the CSTs are shown in pulses per second (pps), and peak EMG is shown equivalent to maximum torque. The bottom plot within each quadrant houses the ensembles, color coordinated in accordance with the color bar, and the CST for reference. The light gray line across plots indicates the time of peak torque. (Ensemble cohorts: N = 240, 215, 202, 232, 232, 229, 210, 259, 202, 107).

### SVR Ensemble Representative Capacity

To highlight the distinct discharge behavior within and across muscles, we display SVR ensemble plots for the TA, MG, and SOL in Figure 7. The visualization capacity and information density inherent to ensembles are easily conveyed here, with distinct differences in discharge characteristics immediately perceptible between muscles.

**Figure 7:**
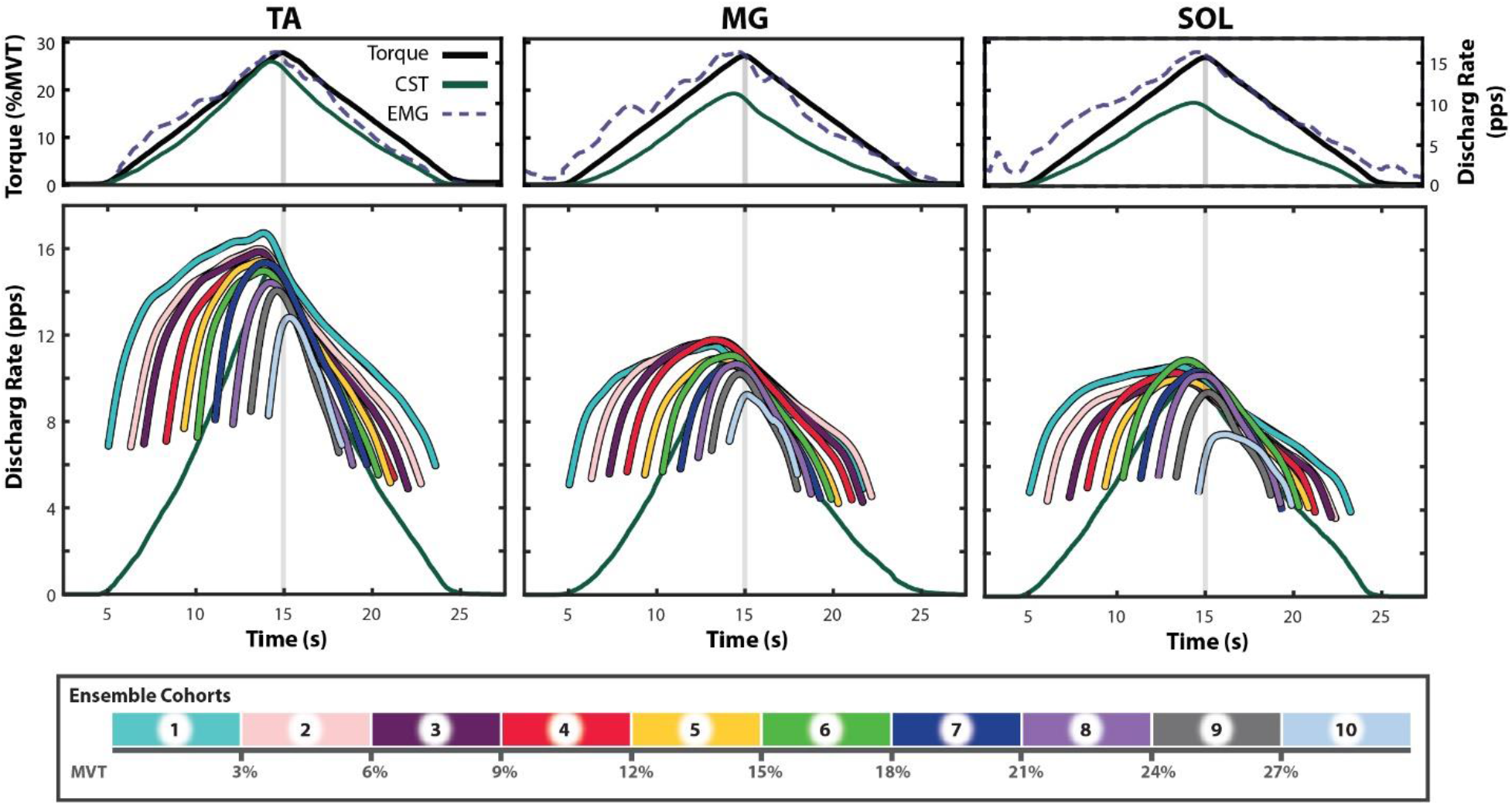
SVR ensembles across muscles. Shown are the ensemble traces for each muscle, generated from motor unit discharge rates estimated with support vector regression (SVR). For each muscle, the top plot depicts the average torque in black, the cumulative spike train (CST) across all units in green, and the average moving root mean squared EMG for all trials in dashed purple. Torque is shown as percent of maximum voluntary torque (MVT), the CSTs are shown in pulses per second (pps), and peak EMG is shown equivalent to maximum torque. The bottom plot for each muscle houses the ensembles, color coordinated in accordance with the color bar, and the CST for reference. The light gray line across plots indicates the time of peak torque. (Ensemble cohorts: [TA: N = 240, 215, 202, 232, 232, 229, 210, 259, 202, 107]; [MG: N = 221, 261, 299, 310, 373, 383, 304, 287, 160, 75]; [SOL: N = 154, 146, 142, 171, 136, 150, 110, 97, 54, 30])

To investigate the ability of the ensembles to portray underlying MU population statistics, we ran a Monte Carlo simulation and show the results in Figure 8. For each iteration of this simulation, we quantified the discharge rate at recruitment, derecruitment, and peak, as well as time to peak discharge rate and ΔF for each ensemble and corresponding population of MUs within that ensemble. To estimate the deviation between ensemble traces and population averages, we fit a linear model to each outcome metric, with fixed effects of ensemble cohort (1-10), whether an estimate was from the random sample or an ensemble trace (method), and the interaction between these two. Results of these linear models are separated by outcome metric as follows.

**Figure 8:**
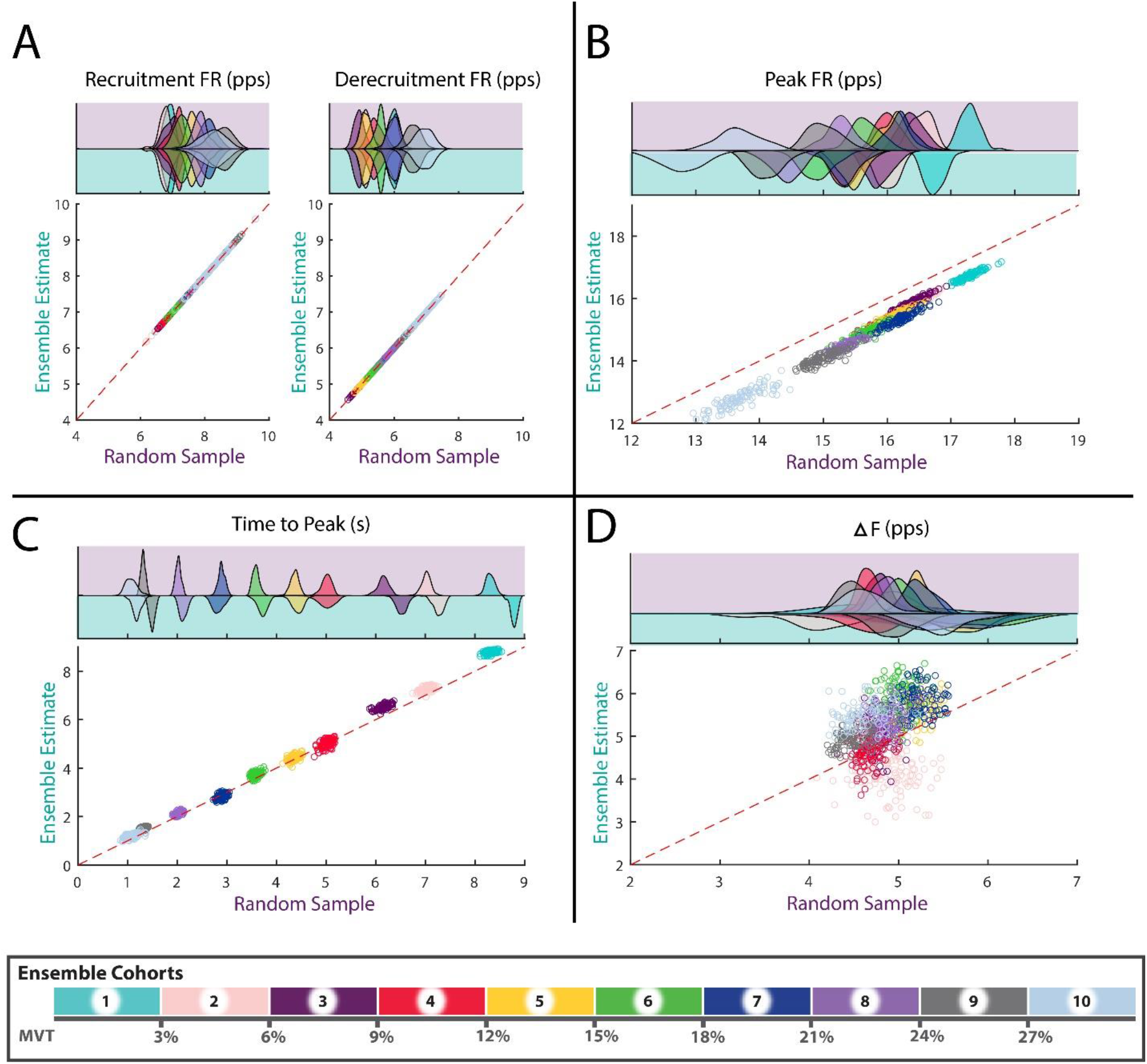
Ensemble accuracy. Shown above are the results of the Monte Carlo simulation, treating the construction of ensembles as a process. This is shown for five parameters of interest, including discharge rate at recruitment (A, left), derecruitment (A, right), and peak (B) as well as time from recruitment to peak discharge rate (C) and ΔF (D). Within each quadrant, the bottom plot depicts the ensemble estimate of a given parameter against the average of that parameter estimate for all individual motor units. This is separated into ensemble cohorts according to the color bar with a data point for each iteration (N = 1000). The dashed red line depicts a theoretical 100% agreement between the ensemble estimates and the population average for a random sample. The probability density for each ensemble and parameter are displayed in the top row of each quadrant and follow an identical color scheme. Distributions outlined in purple on the top row correspond to population averages of the sample with those on the bottom row outlined in blue corresponding to the ensemble estimates. Motor units were sampled from the decomposed population of tibialis anterior units (N = 2128).

#### Peak Discharge Rate

We fit a linear model to peak discharge rate using ordinary least squares (Peak DR ~ Ensemble * Method). The model exhibits substantial explanatory power (R^2^ = 0.96, F(19, 1980) = 2614.2, p < 0.001, adj. R^2^ = 0.96) with significant effects of ensemble (F(9, 1980) = 4851.9, p < 0.001), method (F(1, 1980) = 5762.7, p < 0.001), and their interactions (F(9, 1980) = 26.68, p < 0.001). Averaging across all ensemble cohorts, a main effect comparison of marginal means for method yields an estimated decrease of 0.725 pps (95%CI: [0.706, 0.744]) in the ensemble estimates when compared to the MU population averages. Separating by ensemble cohort, estimates range from a minimum decrease in the ensemble estimates of 0.511 pps (95%CI: [0.452, 0.570]) in ensemble three to a decrease of 0.908 pps (95%CI: [0.849, 0.968]) in ensemble nine.

#### Recruitment & Derecruitment Discharge Rate

We fit a linear model to discharge rate at recruitment and derecruitment, separately, using ordinary least squares (Estimate ~ Ensemble * Method). The models exhibit substantial explanatory power for discharge rate at recruitment (R^2^ = 0.84, F(19, 1980) = 566.2, p < 0.001, adj. R^2^ = 0.84) and derecruitment (R^2^ = 0.92, F(19, 1980) = 1261.6, p < 0.001, adj. R^2^ = 0.92). For both models, the fixed effect of ensemble cohort was significant (recruitment: F(9, 1980) = 1195.4, p < 0.001; derecruitment: F(9, 1980) = 2663.5, p < 0.001), but the method and interaction terms were not.

#### Time to Peak

We fit a linear model to the time from recruitment to peak discharge rate using ordinary least squares (Time to Peak ~ Ensemble * Method). The model exhibits substantial explanatory power (R^2^ = 0.998, F(19, 1980) = 72029.00, p < 0.001, adj. R^2^ = 0.998) with significant effects of ensemble (F(9, 1980) = 151721.00, p < 0.001), method (F(1, 1980) = 1547.77, p < 0.001), and their interactions (F(9, 1980) = 168.25, p < 0.001). Averaging across all ensemble cohorts, a main effect comparison of marginal means for method yields an estimated increase of 0.160 s (95%CI: [0.152, 0.168]) in the ensemble estimates. Separating by ensemble cohort, estimates range from a non-significant effect in ensemble four to a maximum increase of 0.480 pps (95%CI: [0.451, 0.502]) in ensemble one.

#### ΔF

We fit a linear model to the estimates of ΔF for both the ensemble traces and average values within each ensemble (ΔF ~ Ensemble * Method) for ensemble cohorts two through ten, due to a lack of test units for the first ensemble trace. The model exhibits high explanatory power (R^2^ = 0.75, F(17, 1782) = 314.04, p < 0.001, adj. R^2^ = 0.75) with significant effects of ensemble (F(8, 1980) = 382.21, p < 0.001), method (F(1, 1980) = 798.58, p < 0.001), and their interactions (F(8, 1980) = 185.29, p < 0.001). Averaging across all ensemble cohorts, a main effect comparison of marginal means for method yields an estimated increase of 0.361 pps (95%CI: [0.336, 0.386]) in the ensemble estimates. Separating by ensemble cohort, estimates range from a decrease of 0.833 pps (95%CI: [-0.908, −0.758]) in ensemble two, to an increase of 0.890 pps (95%CI: [0.815, 0.965]) in ensemble six.

## DISCUSSION

The increasingly prevalent implementation of HD-sEMG as a research tool has the potential to bolster insights into not only basic human and non-human neurophysiology but both healthy motor control and pathological motor dysfunction. In this paper, we addressed two potential limitations that researchers using HD-sEMG often encounter. That is, 1) the computational approach used to generate smooth discharge rate estimates and 2) the process of visualizing the discharge patterns of large populations of decomposed MUs. To address these potential issues, we suggest support vector regression (SVR) as an improved computational approach over traditionally used methods and put forth the visualization of large populations of MUs as ensembles.

### Estimating Motor Unit Discharge Rate

In its current state, motor neuron and MU research lacks an agreed upon computational method for creating smooth continuous estimates of discharge rates. A recent comprehensive tutorial regarding HD-sEMG expertly detailed the process of extracting neural information from HD-sEMG, including data acquisition, decomposition, an overview of quantifiable MU properties, and motor unit tracking, but did not extend commentary on the optimal fit methods for MU analysis techniques (Del Vecchio et al., 2020). Various computational approaches are routinely employed to generate smooth estimates of discharge rate and have the potential to introduce artifacts in MU quantification and characterization protocols (Hassan et al., 2020). Commonly employed approaches include filtering of decomposed MU spike trains with a window function (i.e. Hanning window) or fitting instantaneous discharge rates with a polynomial function of an arbitrary degree. Though these methods are historically in wide use, investigation of their biasing effects on MU discharge estimates is lacking. In this paper we have highlighted these potential effects and proposed an alternative method, support vector regression, for creating smooth estimates of MU discharge rate.

Support vector regression (SVR), as a computational fitting approach, provides far greater flexibility than Hanning (Hann) window filtering or polynomial regression and allows for greater optimization of fitting characteristics. Specifically, much like the principles of support vector machines (Cortes & Vapnik, 1995; Vapnik, 1995), SVR fits a hyperplane and margin to a collection of training data points, with use of a kernel function to effectively fit the data points in a higher dimensional space. These characteristics, in addition to the ability to independently weight observations, facilitates detailed SVR hyperplane optimization and allows for a collection of data points to be estimated in a highly tunable manner. This is particularly useful in creating smooth estimates of MU discharge rate when compared to traditional approaches, where edge effects from filtering with a window or over smoothing through polynomial regression can perceivable affect data interpretation. Indeed, as can be seen in Figure 3 and Figure 4, SVR appears to offer a suitable middle ground of mitigating noise (i.e. smoothing) while also retaining relevant characteristics of MU discharge (e.g. discharge rate at recruitment, initial amplification, and rate modulation phases).

The ability of SVR to balance the tradeoff between bias and variance was achieved through the tuning of parameters for the kernel and the cost functions and allows SVR to generate more accurate estimates. As shown in Figures 4 and 5, this approach provides more accurate estimates when compared to both filtering with the Hanning window and regression with a polynomial function. When comparing SVR and Hanning window filtering, this is most evident in the fit error produced for the first five MU pulses. In these initial MU pulses, the Hanning fits produced significantly higher deviations across all muscles with an estimated increase in error over SVR fits of 0.758 pps (95%CI: [0.714, 0.802]) for the TA, 0.564 pps (95%CI: 0.521, 0.608]) for the MG, and 0.430 pps (95%CI: [0.363, 0.496]) for the SOL. When Comparing SVR and polynomial regression, the over-smoothing effect of the polynomial fits can be seen in Figure 4 and Figure 5, with the polynomials absolute error higher than the SVR estimates for the first five MU pulses in Figure 5A and entire MU duration in Figure 5B. This increase in absolute error was most severe with the 5^th^ degree polynomial, which produced an average absolute error for the first five instances of discharge that was 0.225 pps (95%CI: [0.195, 0.256]) greater than the error observed with the SVR estimates. This increased error was also apparent across the entire MU duration with an average absolute error at any given instance of 1.052 (95%CI: [1.019, 1.086]) for the 5^th^ degree polynomial, 1.021 pps (95%CI: [0.987, 1.054]) for the 6^th^ degree polynomial, and 0.949 pps (95%CI: [0.915, 0.982]) for the SVR fits. Though the differences between fits is small in magnitude, these values represent the average error at any given instance and will accumulate across the MU duration. The potential effects of these absolute fit errors on choice outcome metrics could be substantial and are apparent in the selected example MU in Figure 3. In this example, the Hanning fit underestimates the initial discharge rate by almost 2 pps and the polynomial functions omit the abrupt decrease in discharge rate on the descending portion of the ramp. The ramifications of these artifacts are blatant in Figure 6, where visualization of the ensembles broadcast the fitting characteristics of each fit method, a key strength of visualizing MU populations as ensembles.

### Population Visualization: Ensembles

The increase in MU yield within studies employing modern HD-sEMG technology necessitates an intuitive approach to displaying the findings of a dataset in an efficient manner. We suggest that large MU datasets be visualized as ensembles. Ensembles are traces of MU discharge rate that represent the average discharge profiles of a subpopulation of motor units and can be used to quickly convey the discharge characteristics of a population of motor units. Here, we have subdivided our MU dataset into ten cohorts based upon torque at recruitment, to observe changes in discharge profile as a function of recruitment threshold, though an alternative parameter could have easily been chosen (e.g. discharge rate at recruitment). Depiction of these ensembles can efficiently show characteristics of the MU populations discharge rate, including changes across the subdividing parameter as well as changes following an intervention should ensembles be constructed for both scenarios.

The capability of ensembles to convey the underlying discharge characteristics of the MU population can be observed in both Figure 2 and Figure 7. In Figure 2A, the ensemble traces appear to overlay the non-normalized SVR fits and represent the average discharge profile of the underlying TA MUs. This is apparent across ensemble cohorts, with the lower cohorts easily represented and the higher ensemble cohorts exhibiting a greater variability in derecruitment that the ensembles capture with the average instance of derecruitment. In Figure 7, distinct modes of MU discharge rate can be observed for each of the ten recruitment threshold cohorts across the TA, MG, and SOL muscles. This occurs in accordance with expected findings and includes an initial acceleration phase, where activation of PICs take place, a gain attenuation phase (post-acceleration rate saturation) where the PICs are likely saturated, a decay in discharge rate with decreases in torque, and subsequent hysteresis where the derecruitment discharge rate is decreased and occurring at lower torque values (Heckman & Enoka, 2012). Interestingly, a semblance of the “onion-skin” phenomena may also be conveyed with these ensemble plots when subdivided by recruitment threshold, of which could be used to further investigate this interesting phenomenon (De Luca & Contessa, 2015; Inglis & Gabriel, 2021; Piotrkiewicz & Türker, 2017). Additionally, the differences between muscles (Figure 7) is stark and highlights the ability of these ensemble traces to efficiently convey differences in discharge rate between groups of MUs. When looking across muscles, the ensembles easily represent muscle specific changes in discharge rate and display comparable trends observed in literature (Kim et al., 2020).

To characterize the relationship between the ensembles and underlying population of MUs, we conducted a simulation where we iteratively resampled our TA MU population, taking two-thirds of the population each time to generate ensembles. Results of this simulation illuminate the rich density of information garnered through the ensemble traces. Of note, the ensemble traces appear to represent the recruitment and derecruitment discharge rate nearly identically (Figure 8A), a key feature woven into their construction, while systemically under-estimating the peak discharge rate (Figure 8B). This under-estimation can likely be attributed to the aligning process, where instances of recruitment and derecruitment are aligned rather than the instance of peak discharge. Though the instances of peak discharge rate across MUs within a given ensemble are likely similar, there is a slight smoothing effect as peaks are misaligned. Across all ensembles this is estimated as an approximate attenuation of 0.725 pps (95%CI: [0.706, 0.744]). Furthermore, though the time to peak discharge rate of the ensemble traces appear to closely track that of the MU population (Figure 8C), they are not identical. Across all ensembles, a delay of 0.160 s (95%CI: [0.152, 0.168]) is estimated and should be considered when employing ensemble traces for quantification. Similarly, considering the quantification of ΔF across the population or through the ensemble traces, the ensembles produce estimates approximately 0.361 pps (95%CI: [0.336, 0.386]) higher across all ensembles (Figure 8D).

Though the ensemble estimates of peak discharge rate, time to this peak, and ΔF are significantly different than the underlying MU population tested here, their magnitudes and intended purpose must be considered. Explicitly, these deviations must be interpreted with the understanding that the ensembles are designed as a visualization tool to supplement the proper analysis and reporting of population statistics. With the proper analysis and reporting of population statistics, ensemble traces can be employed to quickly convey changes in discharge profile across the population with a level of accuracy sufficient for visualization purposes. Indeed, depending on scale, qualitative considerations such as line thickness can easily obscure the visualization of time to peak discharge by more than 160 ms. With this in mind, visualization of ensembles allow for researchers to supplement their quantitative findings and portray MU discharge characteristics of an entire dataset in an intuitive manner where the various modes of MU discharge are observed.

### Further Considerations

On a fundamental level, the visualization of ensemble traces are meant to portray average discharge characteristics of MU populations as an input-output function. Here we have chosen an input of torque at recruitment and isolated ten cohorts of MUs, though this can conceivably be expanded to a variety of neuronal discharge recordings and types of input variables. For example, the discharge patterns of cortical neurons within the auditory cortex could be separated by the frequency of auditory stimulation and visualized in a similar manner to what was conducted here (Bitterman, Mukamel, Malach, Fried, & Nelken, 2008; Montgomery & Wehr, 2010). Additionally, the modulation of biceps brachii MUs during isometric elbow flexion ramps, subdivided by various degrees of deltoid activation, in individuals with chronic hemiparetic stroke could be used in a similar fashion to construct ensemble traces. This would supply unique insight into the prevalent coupling of shoulder abduction with elbow, wrist, and finger flexion post stroke (Dewald & Beer, 2001; Dewald, Pope, Given, Buchanan, & Rymer, 1995). More closely related to the paper at hand, we could have separated MUs into groups of equal sample sizes based on their torque at recruitment and gauged insight into recruitment spacing at the cost of insight into distinct differences across recruitment threshold.

### Potential Limitations

Though the methods outlined here represent an incremental step in the analysis and visualization of MU discharge profiles, a few limitations must be considered. In specific, the biases introduced by each fit method highlighted here are only relevant to quantitative outcome metrics that employ smooth estimates of MU discharge. The benefits of any computational approach are dependent on their application. Metrics that analyze adjacent aspects of MU discharge (e.g. inter-spike interval) would remain unaffected by the various fitting schemes. Furthermore, the improvement in estimates achieved through SVR are only improvements upon the two most commonly employed methods used in fitting spinal MU discharge rates. We are not proposing that SVR is the ultimate fitting method, that alternative approaches are inferior, or that improvements to the current approaches (e.g. bias correction for the windowing) are unavailable. Instead, we are emphasizing SVR as a modern approach that outperforms commonly employed methods. Of particular note, though SVR outperforms a 6^th^ degree polynomial, implementation of a 6^th^ degree polynomial appears to outperform the commonly employed 5^th^ degree polynomial, and thus could be considered in various circumstances. Lastly, more extensive hyperparameter optimization methods, such as a Bayesian optimization scheme, may provide superior results.

## CONCLUSIONS

To address potential limitations in the analysis and visualization of large MU populations in modern HD-sEMG studies, smooth estimates of MU discharge rates can be generated with SVR and MU populations may be visualized as ensembles. In this study, we have shown SVR to be an effective computational procedure for generating smooth estimates of MU discharge rate. In addition to possessing superior adaptability, when compared to Hanning window filtering and polynomial regression, SVR more accurately estimates the recruitment region of MU discharge while maintaining adequate accuracy throughout the duration of discharge. These desirable characteristics of SVR are highly evident when used to generate ensembles. The generation of ensembles, as defined in this study, represents a novel method to visualize the average discharge profiles of many MUs within a dataset. This allows for the efficient rendering of discharge characteristics that are representative of the entire dataset and not comprised of single choice example trials or a barrage of scatter plots. In combination, the use of SVR and generation of ensembles represents an efficient approach for portraying population discharge characteristics with appropriate accuracy for effective visualization.

## REFERENCES

Afsharipour, B., Manzur, N., Duchcherer, J., Fenrich, K. F., Thompson, C. K., Negro, F.,. … Gorassini, M. A. (2020). Estimation of self-sustained activity produced by persistent inward currents using firing rate profiles of multiple motor units in humans. Journal of neurophysiology, 124(1), 63–85. doi:10.1152/jn.00194.2020

Bennett, D. J., Li, Y., Harvey, P. J., & Gorassini, M. (2001). Evidence for plateau potentials in tail motoneurons of awake chronic spinal rats with spasticity. Journal of neurophysiology, 86(4), 1972–1982. doi:10.1152/jn.2001.86.4.1972

Bitterman, Y., Mukamel, R., Malach, R., Fried, I., & Nelken, I. (2008). Ultra-fine frequency tuning revealed in single neurons of human auditory cortex. Nature, 451(7175), 197–201. doi:10.1038/nature06476

Boccia, G., Martinez-Valdes, E., Negro, F., Rainoldi, A., & Falla, D. (2019). Motor unit discharge rate and the estimated synaptic input to the vasti muscles is higher in open compared with closed kinetic chain exercise. J Appl Physiol (1985), 127(4), 950–958. doi:10.1152/japplphysiol.00310.2019

Cortes, C., & Vapnik, V. (1995). Support-vector networks. Machine Learning, 20(3), 273–297. doi:10.1007/bf00994018

Cristianini, N., & Shawe-Taylor, J. (2000). An introduction to support vector machines and other kernel-based learning methods: Cambridge university press.

De Luca, C. J., Adam, A., Wotiz, R., Gilmore, L. D., & Nawab, S. H. (2006). Decomposition of surface EMG signals. Journal of neurophysiology, 96(3), 1646–1657. doi:10.1152/jn.00009.2006

De Luca, C. J., & Contessa, P. (2015). Biomechanical benefits of the Onion-Skin motor unit control scheme. Journal of Biomechanics, 48(2), 195–203. doi:10.1016/j.jbiomech.2014.12.003

De Luca, C. J., LeFever, R. S., McCue, M. P., & Xenakis, A. P. (1982a). Behaviour of human motor units in different muscles during linearly varying contractions. J Physiol, 329, 113–128. doi:10.1113/jphysiol.1982.sp014293

De Luca, C. J., LeFever, R. S., McCue, M. P., & Xenakis, A. P. (1982b). Behaviour of human motor units in different muscles during linearly varying contractions. The Journal of Physiology, 329, 113–128. doi:10.1113/jphysiol.1982.sp014293

Del Vecchio, A., Holobar, A., Falla, D., Felici, F., Enoka, R., & Farina, D. (2020). Tutorial: Analysis of motor unit discharge characteristics from high-density surface EMG signals. Journal of Electromyography and Kinesiology, 102426.

Dewald, J. P., & Beer, R. F. (2001). Abnormal joint torque patterns in the paretic upper limb of subjects with hemiparesis. Muscle & nerve, 24(2), 273–283. doi:10.1002/1097-4598(200102)24:2<273∷aid-mus130>3.0.co;2-z

Dewald, J. P., Pope, P. S., Given, J. D., Buchanan, T. S., & Rymer, W. Z. (1995). Abnormal muscle coactivation patterns during isometric torque generation at the elbow and shoulder in hemiparetic subjects. Brain: a journal of neurology, 118 (Pt 2, 495–510. doi:DOI 10.1093/brain/118.2.495

Drucker, H., Burges, C., Kaufman, L., Smola, A., & Vapnik, V. N. (1996). Support Vector Regression Machines. Paper presented at the Advances in Neural Information Processing Systems 9: Proceedings of the 1996 Conference.

Farina, D., Holobar, A., Gazzoni, M., Zazula, D., Merletti, R., & Enoka, R. M. (2009). Adjustments differ among low-threshold motor units during intermittent, isometric contractions. Journal of neurophysiology, 101(1), 350–359. doi:10.1152/jn.90968.2008

Farina, D., Holobar, A., Merletti, R., & Enoka, R. M. (2010). Decoding the neural drive to muscles from the surface electromyogram. Clin Neurophysiol, 121(10), 1616–1623. doi:10.1016/j.clinph.2009.10.040

Gorassini, M., Yang, J. F., Siu, M., & Bennett, D. J. (2002). Intrinsic Activation of Human Motoneurons: Possible Contribution to Motor Unit Excitation. Journal of neurophysiology, 87(4), 1850–1858. doi:10.1152/jn.00024.2001

Hassan, A., Thompson, C. K., Negro, F., Cummings, M., Powers, R. K., Heckman, C. J.,. .. McPherson, L. M. (2020). Impact of parameter selection on estimates of motoneuron excitability using paired motor unit analysis. Journal of Neural Engineering, 17(1), 016063. doi:10.1088/1741-2552/ab5eda PMID - 31801123

Heckman, C. J., & Enoka, R. M. (2012). Motor unit. Comprehensive Physiology, 2(4), 2629–2682. doi:10.1002/cphy.c100087

Holobar, A., Minetto, M. A., & Farina, D. (2014). Accurate identification of motor unit discharge patterns from high-density surface EMG and validation with a novel signal-based performance metric. J Neural Eng, 11(1), 016008. doi:10.1088/1741-2560/11/1/016008

Hug, F., Avrillon, S., Del Vecchio, A., Casolo, A., Ibanez, J., Nuccio, S.,. .. Farina, D. (2021). Analysis of motor unit spike trains estimated from high-density surface electromyography is highly reliable across operators. J Electromyogr Kinesiol, 58, 102548. doi:10.1016/j.jelekin.2021.102548

Inglis, J. G., & Gabriel, D. A. (2021). Is the ‘reverse onion skin’ phenomenon more prevalent than we thought during intramuscular myoelectric recordings from low to maximal force outputs? Neuroscience letters, 743, 135583. doi:https://doi.org/10.1016/j.neulet.2020.135583

Johnson, M. D., Thompson, C. K., Tysseling, V. M., Powers, R. K., & Heckman, C. J. (2017). The potential for understanding the synaptic organization of human motor commands via the firing patterns of motoneurons. Journal of neurophysiology, 118(1), 520–531. doi:10.1152/jn.00018.2017

Kallenberg, L. A., & Hermens, H. J. (2009). Motor unit properties of biceps brachii in chronic stroke patients assessed with high-density surface EMG. Muscle Nerve, 39(2), 177–185. doi:10.1002/mus.21090

Kim, E. H., Wilson, J. M., Thompson, C. K., & Heckman, C. J. (2020). Differences in estimated persistent inward currents between ankle flexors and extensors in humans. Journal of neurophysiology, 124(2), 525–535. doi:10.1152/jn.00746.2019

Li, X., Holobar, A., Gazzoni, M., Merletti, R., Rymer, W. Z., & Zhou, P. (2015). Examination of Poststroke Alteration in Motor Unit Firing Behavior Using High-Density Surface EMG Decomposition. IEEE Trans Biomed Eng, 62(5), 1242–1252. doi:10.1109/TBME.2014.2368514

Martinez-Valdes, E., Negro, F., Laine, C. M., Falla, D., Mayer, F., & Farina, D. (2017). Tracking motor units longitudinally across experimental sessions with high-density surface electromyography. J Physiol, 595(5), 1479–1496. doi:10.1113/JP273662

Montgomery, N., & Wehr, M. (2010). Auditory cortical neurons convey maximal stimulus-specific information at their best frequency. The Journal of Neuroscience, 30(40), 13362–13366. doi:10.1523/JNEUROSCI.2899-10.2010

Murphy, S. A., Negro, F., Farina, D., Onushko, T., Durand, M., Hunter, S. K.,. .. Hyngstrom, A. (2018). Stroke increases ischemia-related decreases in motor unit discharge rates. Journal of neurophysiology, 120(6), 3246–3256. doi:10.1152/jn.00923.2017

Nawab, S. H., Chang, S. S., & De Luca, C. J. (2010). High-yield decomposition of surface EMG signals. Clin Neurophysiol, 121(10), 1602–1615. doi:10.1016/j.clinph.2009.11.092

Negro, F., & Farina, D. (2012). Factors influencing the estimates of correlation between motor unit activities in humans. PLoS One, 7(9), e44894. doi:10.1371/journal.pone.0044894

Negro, F., Muceli, S., Castronovo, A. M., Holobar, A., & Farina, D. (2016). Multi-channel intramuscular and surface EMG decomposition by convolutive blind source separation. Journal of Neural Engineering, 13(2), 026027.

Orssatto, L. B. R., Mackay, K., Shield, A. J., Sakugawa, R. L., Blazevich, A. J., & Trajano, G. S. (2021). Estimates of persistent inward currents increase with the level of voluntary drive in low-threshold motor units of plantar flexor muscles. Journal of neurophysiology, 125(5), 1746–1754. doi:10.1152/jn.00697.2020

Oya, T., Riek, S., & Cresswell, A. G. (2009). Recruitment and rate coding organisation for soleus motor units across entire range of voluntary isometric plantar flexions. J Physiol, 587(Pt 19), 4737–4748. doi:10.1113/jphysiol.2009.175695

Piotrkiewicz, M., & Türker, K. S. (2017). Onion Skin or Common Drive? Frontiers in cellular neuroscience, 11, 2–2. doi:10.3389/fncel.2017.00002

Powers, R. K., Nardelli, P., & Cope, T. C. (2008). Estimation of the contribution of intrinsic currents to motoneuron firing based on paired motoneuron discharge records in the decerebrate cat. Journal of neurophysiology, 100(1), 292–303. doi:10.1152/jn.90296.2008 PMID – 18463182

Rau, G., & Disselhorst-Klug, C. (1997). Principles of high-spatial-resolution surface EMG (HSR-EMG): single motor unit detection and application in the diagnosis of neuromuscular disorders. Journal of Electromyography and Kinesiology, 7(4), 233–239. doi:10.1016/s1050-6411(97)00007-2

Smola, A. J., & Schölkopf, B. (1998). Learning with kernels (Vol. 4): Citeseer.

Smola, A. J., & Schölkopf, B. (2004). A tutorial on support vector regression. Statistics and Computing, 14(3), 199–222. doi:10.1023/b:Stco.0000035301.49549.88

Stephenson, J. L., & Maluf, K. S. (2011). Dependence of the paired motor unit analysis on motor unit discharge characteristics in the human tibialis anterior muscle. Journal of Neuroscience Methods, 198(1), 84–92. doi:10.1016/j.jneumeth.2011.03.018

Udina, E., D’Amico, J., Bergquist, A. J., & Gorassini, M. A. (2010). Amphetamine increases persistent inward currents in human motoneurons estimated from paired motor-unit activity. Journal of neurophysiology, 103(3), 1295–1303. doi:10.1152/jn.00734.2009 PMID - 20053846

Vapnik, V. N. (1995). The Nature of Statistical Learning Theory: Springer.

Wilson, J. M., Thompson, C. K., Miller, L. C., & Heckman, C. J. (2015). Intrinsic excitability of human motoneurons in biceps brachii versus triceps brachii. Journal of neurophysiology, 113(10), 3692–3699. doi:10.1152/jn.00960.2014

